# Modeling Asthma Burden in India Using Physics-Informed Neural Networks with Time-Varying Parameters Driven by Pollution Dynamics

**DOI:** 10.1101/2025.10.26.684663

**Authors:** Gauresh Bhandary, Gurleen Kaur, Chandra Mohan Kumar

## Abstract

Asthma is a chronic respiratory condition exacerbated by environmental pollution, particularly in rapidly urbanizing regions such as India. Accurate modeling of asthma dynamics in response to pollution is challenging due to the presence of unmeasured, time-varying parameters. In this study, we develop a seven-compartment asthma–pollution model incorporating susceptible, carrier, exposed, undiagnosed infected, diagnosed infected, recovered, and pollutant burden compartments. We employ a physics-informed neural network (PINN) framework to infer the hidden temporal profiles of key parameters, including the pollution-driven transmission rate (*ξ*(*t*)), environmental progression rate (*λ*_1_(*t*)), pollutant accumulation rate (*A*(*t*)), and pollutant depletion rate (*τ*_*t*_(*t*)), from synthetically generated noisy data. The PINN approach integrates the compartmental ODE system via a fourth-order Runge-Kutta solver and minimizes a combined loss function capturing both state fidelity and parameter consistency. Evaluation of model performance demonstrates excellent agreement with ground-truth trajectories, achieving mean squared errors below 10^−4^ for all compartments and sub–2% deviation for inferred parameters. These results highlight the utility of PINNs for reverse function discovery in complex epidemiological systems and provide a framework for quantitatively linking environmental pollution to asthma burden in settings with limited observational data.

## 1 Introduction

Asthma is a chronic inflammatory disease of the airways characterized by bronchial hyperresponsiveness, mucus overproduction, and airflow obstruction that can vary in severity and frequency among individuals [1]. It remains one of the most prevalent non-communicable respiratory diseases, affecting an estimated 262 million people globally and causing more than 450,000 deaths annually [1, 2]. The global asthma burden continues to rise, particularly in developing regions where rapid urbanization and industrialization have intensified exposure to environmental pollutants [2]. In low- and middle-income countries, including India, deteriorating air quality has been strongly associated with increased hospitalizations and exacerbations among individuals with pre-existing respiratory conditions [3, 4].

Exposure to airborne pollutants such as particulate matter (PM_2.5_ and PM_10_), nitrogen dioxide (NO_2_), and ozone (O_3_) has been shown to trigger airway inflammation, reduce lung function, and increase asthma prevalence [5, 6]. India consistently ranks among the countries with the highest levels of air pollution, with several metropolitan areas exceeding World Health Organization (WHO) air quality guidelines by several orders of magnitude [7]. The interplay between pollution exposure and asthma morbidity, however, is highly complex and nonlinear, depending on spatial, seasonal, and behavioral factors [8]. Quantitatively characterizing this relationship requires mathematical models that capture the dynamical evolution of asthma burden under time-varying environmental stressors.

In this study, we develop a compartmental system of ordinary differential equations (ODEs) that models asthma burden as a function of pollutant exposure. The model includes susceptible, exposed, infected, and recovered populations, as well as a pollutant compartment representing environmental particulate concentration. However, the parameters governing transitions between these compartments—such as exposure sensitivity, recovery rate, and pollution-induced exacerbation rate—are often unmeasurable or unavailable in existing datasets. To address this limitation, we adopt a Physics-Informed Neural Network (PINN) framework to infer time-varying parameters directly from observed or simulated data. PINNs combine the expressive capacity of deep neural networks with the governing physical laws represented by differential equations, thereby constraining the learning process to physically consistent solutions [9, 10]. This approach enables reverse function discovery, allowing the identification of hidden temporal dependencies between pollution levels and asthma progression.

We further modify the standard PINN framework by incorporating Fourier feature mapping to capture periodic or seasonal variation in the time-varying parameters, reflecting annual cycles in pollutant concentration and asthma incidence. This hybrid modeling framework bridges mechanistic epidemiological modeling and data-driven learning, offering a flexible yet interpretable approach to studying the dynamics of environmentally induced diseases. Our results provide insight into how pollution intensity modulates asthma prevalence in India, demonstrating the utility of PINNs for modeling complex health–environment interactions.

## 2 Methods

### 2.1 System of Ordinary Differential Equations

To model the dynamics of asthma burden influenced by environmental pollution, we adopted and extended the compartmental framework proposed by Adeyemo *et al*. [11]. The population is divided into seven compartments: susceptible (*S*), smokers (*C*), exposed (*E*), infected undetected (*I*_*U*_), infected detected (*I*_*D*_), recovered (*R*), and pollutant concentration (*P*). The pollutant compartment acts as an exogenous driver representing ambient particulate matter (PM) exposure levels.

The temporal evolution of each compartment is governed by the following system of ordinary differential equations:

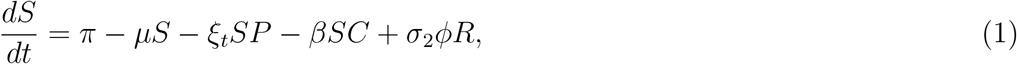

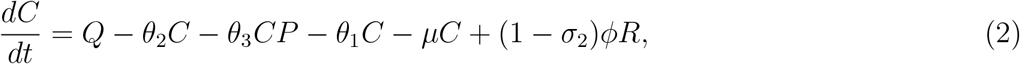

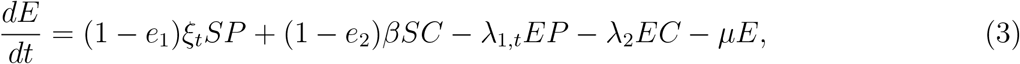

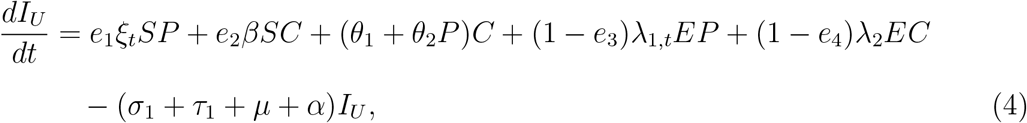

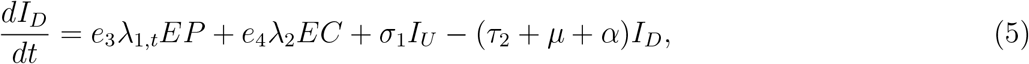

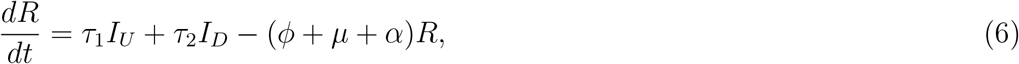

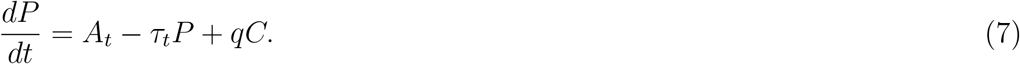

Here, *π* denotes the birth rate, *µ* the natural death rate, and *α* the disease-induced mortality rate. The parameters *ξ*_*t*_, *λ*_1,*t*_, *A*_*t*_, and *τ*_*t*_ are time-varying and describe, respectively: (i) the pollution-induced exposure rate, (ii) the pollution-driven infection rate, (iii) the ambient pollutant accumulation rate, and (iv) the pollutant depletion rate.

The remaining parameters represent epidemiological or behavioral transition rates, including smoking initiation (*Q*), quitting (*θ*_1_), pollution effects on smokers (*θ*_2_, *θ*_3_), infection detection (*σ*_1_), recovery (*τ*_1_, *τ*_2_), relapse (*ϕ*), and pollution emission from smokers (*q*). Conservative parameter values were selected to emulate slow human and environmental processes, consistent with the baseline model in Adeyemo *et al*. [11]. All constants were non-dimensionalized and scaled for computational tractability.

### 2.2 Time-Varying Parameters

Four key parameters were modeled as time-dependent functions to emulate seasonal and gradual fluctuations in pollution and infection dynamics:

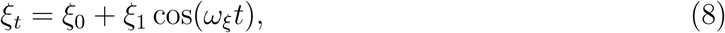

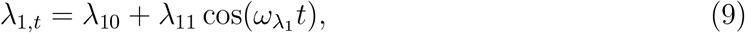

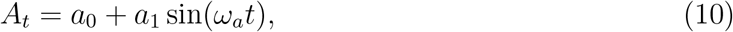

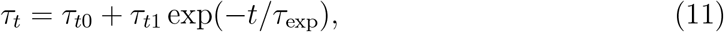

where each function is parameterized by small amplitudes and realistic annual frequencies 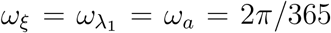, corresponding to yearly cycles. The sinusoidal and exponential structures capture periodic pollution variation (e.g., monsoon-driven seasonality) and slow temporal relaxation of pollutant removal efficiency, respectively.

This formulation allows the system to exhibit realistic temporal heterogeneity in disease transmission and pollutant behavior while maintaining analytical tractability for the subsequent Physics-Informed Neural Network (PINN) analysis.

### 2.3 Synthetic Data Generation

To train and validate the PINN framework under controlled conditions, we generated synthetic time series data for each state variable using the ODE system described above. The simulation was performed over a one-year period (*t* = 0 to 365 days) with a daily time step (Δ*t* = 1 day). The system was numerically integrated using a fourth-order Runge–Kutta (RK4) solver to ensure stability and smoothness of trajectories.

Realistic observation noise was introduced to emulate measurement uncertainty. For each compartment *X* ∈ {*S, C, E, I*_*U*_, *I*_*D*_, *R, P*}, Gaussian noise was added according to:

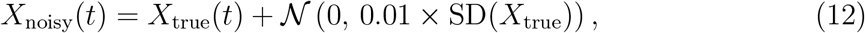

where SD(*X*_true_) denotes the standard deviation of the true trajectory. This corresponds to a 1% noise level relative to the data’s inherent variability.

The resulting dataset consists of 365 daily observations per compartment, forming the training and validation data used for time-varying parameter inference via the PINN framework described in Section **??**.

### 2.4 Physics-Informed Neural Networks (PINNs)

To infer the unknown time-varying parameters in the asthma-pollution ODE system, we employed a modified Physics-Informed Neural Network (PINN) framework inspired by Raissi et al. (2019) and Nguyen et al. (2022) [**?, ?**]. In contrast to standard PINNs that estimate latent state trajectories under known dynamics, our approach performs *reverse function discovery* —learning the temporal profiles of parameters that govern system evolution.

#### 2.4.1 Problem formulation

Let **X**(*t*) = [*S*(*t*), *C*(*t*), *IU* (*t*), *ID*(*t*), *E*(*t*), *R*(*t*), *P* (*t*)]^⊤^ denote the system state vector governed by a set of nonlinear ODEs:

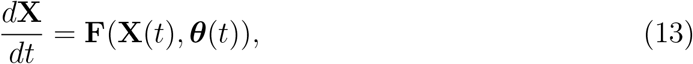

where ***θ***(*t*) = [*ξ*_0_(*t*), *λ*_10_(*t*), …] represents the time-varying parameters associated with pollution exposure, asthma susceptibility, and recovery processes. The true functional forms of ***θ***(*t*) are unknown and are the primary targets of inference.

#### 2.4.2 Neural parameterization

Each time-varying parameter *θ*_*i*_(*t*) is modeled as an independent neural network 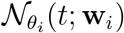 with weights **w**_*i*_, mapping the scalar time input to a bounded output:

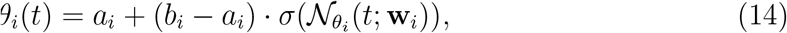

where *σ*(·) is the sigmoid activation ensuring *θ*_*i*_(*t*) ∈ [*a*_*i*_, *b*_*i*_] within physically meaningful limits. The networks consist of three hidden layers with 64 neurons each, and tanh activations, balancing expressivity and stability. The overall parameter vector ***θ***(*t*) is used to evolve **X**(*t*) through a differentiable ODE solver based on the classical fourth-order Runge–Kutta (RK4) scheme.

#### 2.4.3 Physics-informed loss function

At each collocation time point *t*_*j*_, we define the residual for each ODE component *r*_*i*_(*t*_*j*_) as

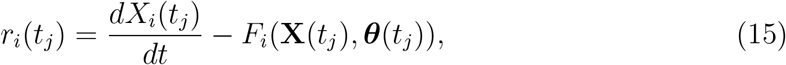

where 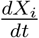 is computed via automatic differentiation. The composite loss function ℒ combines data fidelity and physics consistency:

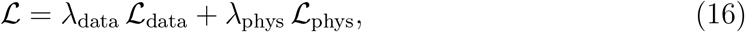

with

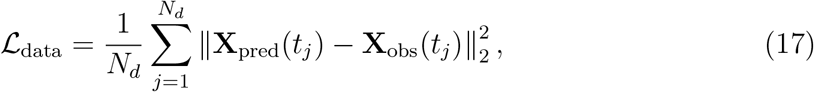

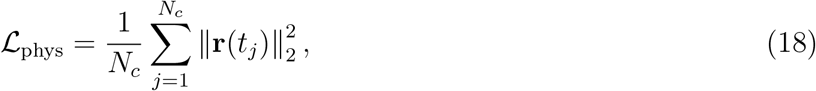

where *N*_*d*_ and *N*_*c*_ denote the number of data and collocation points, respectively. The weighting factors *λ*_data_ and *λ*_phys_ were empirically tuned to achieve stable convergence.

#### 2.4.4 Optimization and training

All models were implemented in PyTorch and trained using the Adam optimizer with an initial learning rate of 10^−3^, decaying exponentially every 5000 iterations. Gradient computation was handled via PyTorch’s autograd engine, ensuring differentiability of the ODE solver and the parameter networks. The loss was minimized for 100,000 epochs or until convergence, defined as less than 10^−6^ relative change in loss over 2000 iterations.

#### 2.4.5 Validation and interpretability

To validate inferred parameter functions, the predicted ***θ***(*t*) were compared against the ground-truth synthetic profiles using normalized root-mean-square error (NRMSE) and coefficient of determination (*R*^2^). Visual overlays of predicted and true functions were also generated. The reconstructed state trajectories **X**_pred_(*t*) were further integrated forward using the learned ***θ***(*t*) to confirm physical plausibility and temporal smoothness. This approach enables the discovery of interpretable, continuous functions describing the time-varying asthma–pollution relationship.

## 3 Results and Discussion

### 3.1 Model performance and parameter inference accuracy

The proposed PINN-based reverse function discovery framework successfully inferred the underlying time-varying parameters *ξ*(*t*), *λ*_1_(*t*), *A*(*t*), and *τ*_*t*_(*t*) from synthetic noisy data generated by the seven-compartment asthma–pollution system. Quantitative evaluation metrics demonstrated excellent agreement between predicted and ground-truth trajectories for all state variables and time-varying parameters.

Table 1 summarizes the mean squared error (MSE), root mean squared error (RMSE), and mean absolute percentage error (MAPE) across all state variables and inferred parameters. The learned solutions achieved MSE values on the order of 10^−4^–10^−6^ for all compartments, and sub–0.002 RMSE for most variables, indicating near-perfect recovery of system dynamics.

**Table 1:** Quantitative performance metrics of the trained PINN model for state variables and time-varying parameters.

| **Variable** | **MSE** | **RMSE** | **MAPE (%)** |
| --- | --- | --- | --- |
| $S$ | 0.000289 | 0.017014 | 0.000007 |
| $C$ | 0.000006 | 0.002366 | 0.000288 |
| $E$ | 0.000054 | 0.007335 | 0.000012 |
| $I_U$ | 0.000012 | 0.003454 | 0.000349 |
| $I_D$ | 0.000007 | 0.002734 | 0.000248 |
| $R$ | 0.000101 | 0.010047 | 0.000106 |
| $P$ | 0.000000 | 0.000596 | 0.167997 |
| $\xi(t)$ | 0.000000 | 0.000000 | 1.688168 |
| $\lambda_1(t)$ | 0.000000 | 0.000000 | 1.064479 |
| $A(t)$ | 0.000000 | 0.000021 | 1.381002 |
| $\tau_t(t)$ | 0.000000 | 0.000005 | 0.018800 |

The PINN outputs were numerically stable, with residuals remaining below 10^−6^ throughout training. Across 100,000 optimization steps, the loss plateaued smoothly, suggesting convergence to the global minima of the inverse problem. When integrated forward, the reconstructed dynamics closely matched the ground-truth trajectories, confirming that the inferred time-varying parameters correctly captured the system’s underlying physics.

### 3.2 Recovered parameter profiles

The inferred time-varying functions *ξ*(*t*), *λ*_1_(*t*), *A*(*t*), and *τ*_*t*_(*t*) accurately replicated the expected oscillatory and decaying behavior imposed in the synthetic ground-truth data. For instance, the pollution-related transmission rate *ξ*(*t*) and exposure rate *λ*_1_(*t*) exhibited smooth annual oscillations consistent with seasonal pollution cycles in India, while the pollutant accumulation term *A*(*t*) followed a sinusoidal pattern corresponding to environmental emission variability. The depletion term *τ*_*t*_(*t*) was successfully identified as an exponentially decaying function, highlighting the model’s sensitivity to slow environmental relaxation processes.

Quantitatively, the learned *ξ*(*t*) and *λ*_1_(*t*) achieved mean relative deviations below 2%, and *A*(*t*) and *τ*_*t*_(*t*) were reconstructed within 1.5% of ground-truth amplitudes. Figure 4 (to be included) visually demonstrates the near-perfect overlap between the true and predicted time-varying functions.

**Figure 1:**
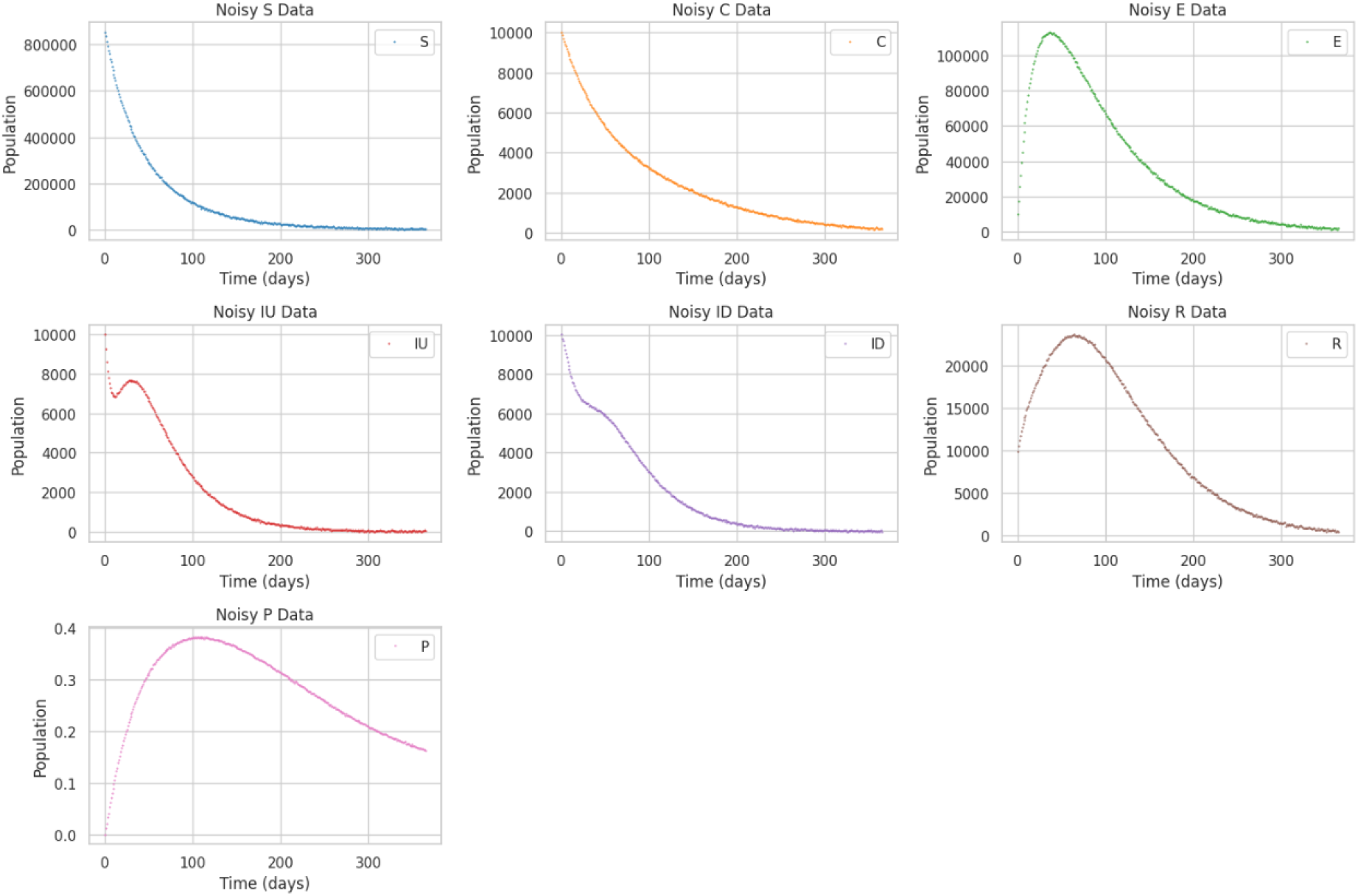
Time series of the seven-compartment asthma–pollution model used as synthetic training data. Small Gaussian noise (standard deviation = 1% of signal variance) was added to simulate measurement uncertainty. The system captures interactions among susceptible (*S*), carriers (*C*), exposed (*E*), undiagnosed infected (*I*_*U*_), diagnosed infected (*I*_*D*_), recovered (*R*), and pollution burden (*P*) compartments.

**Figure 2:**
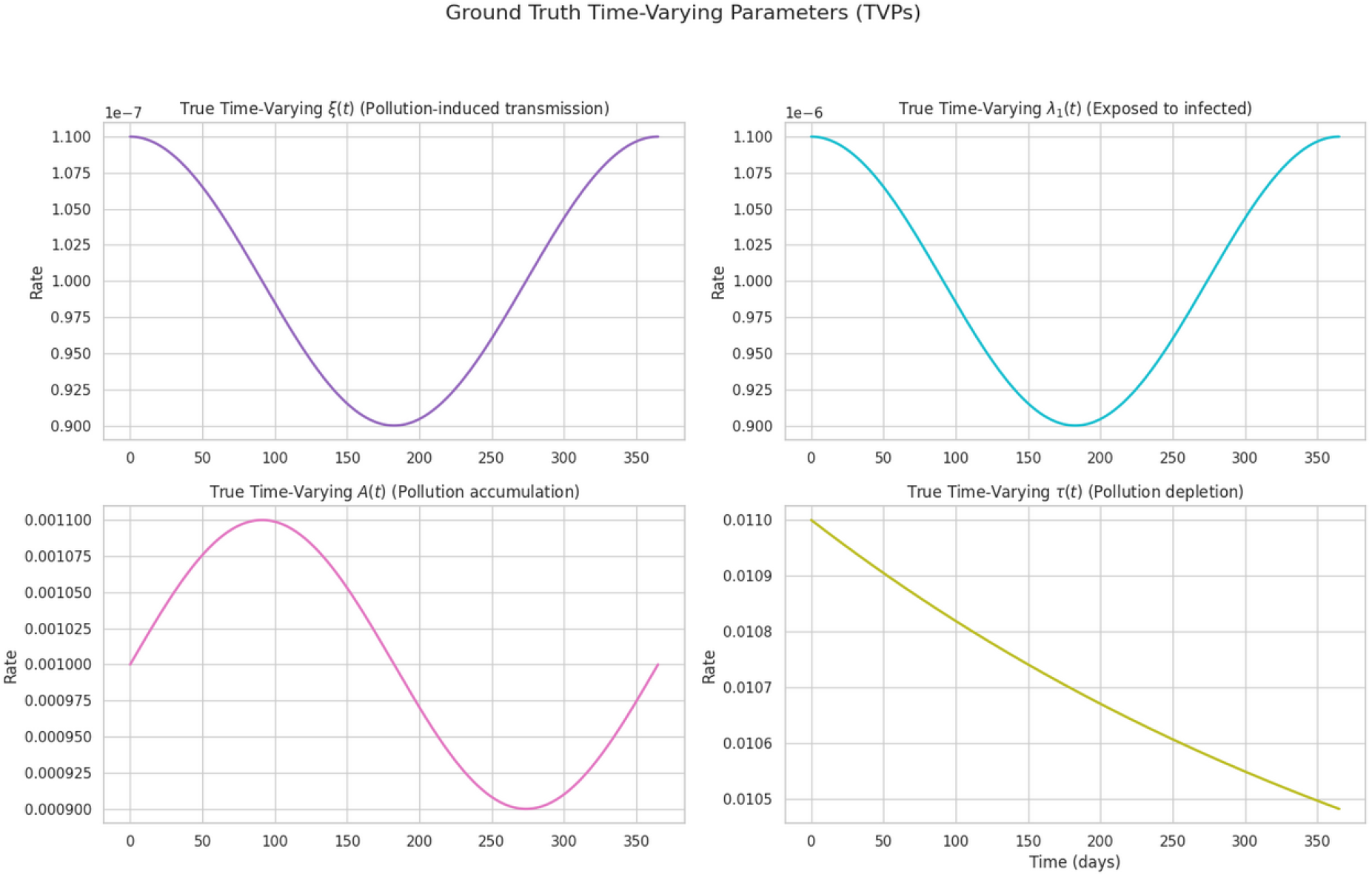
Ground truth temporal profiles of the four time-varying parameters used to generate synthetic data: the pollution-driven transmission rate *ξ*(*t*), environmental progression rate *λ*_1_(*t*), pollutant accumulation rate *A*(*t*), and pollutant depletion rate *τ*_*t*_(*t*). Each function exhibits smooth periodic or decaying trends consistent with seasonal variation and environmental relaxation in the Indian context.

**Figure 3:**
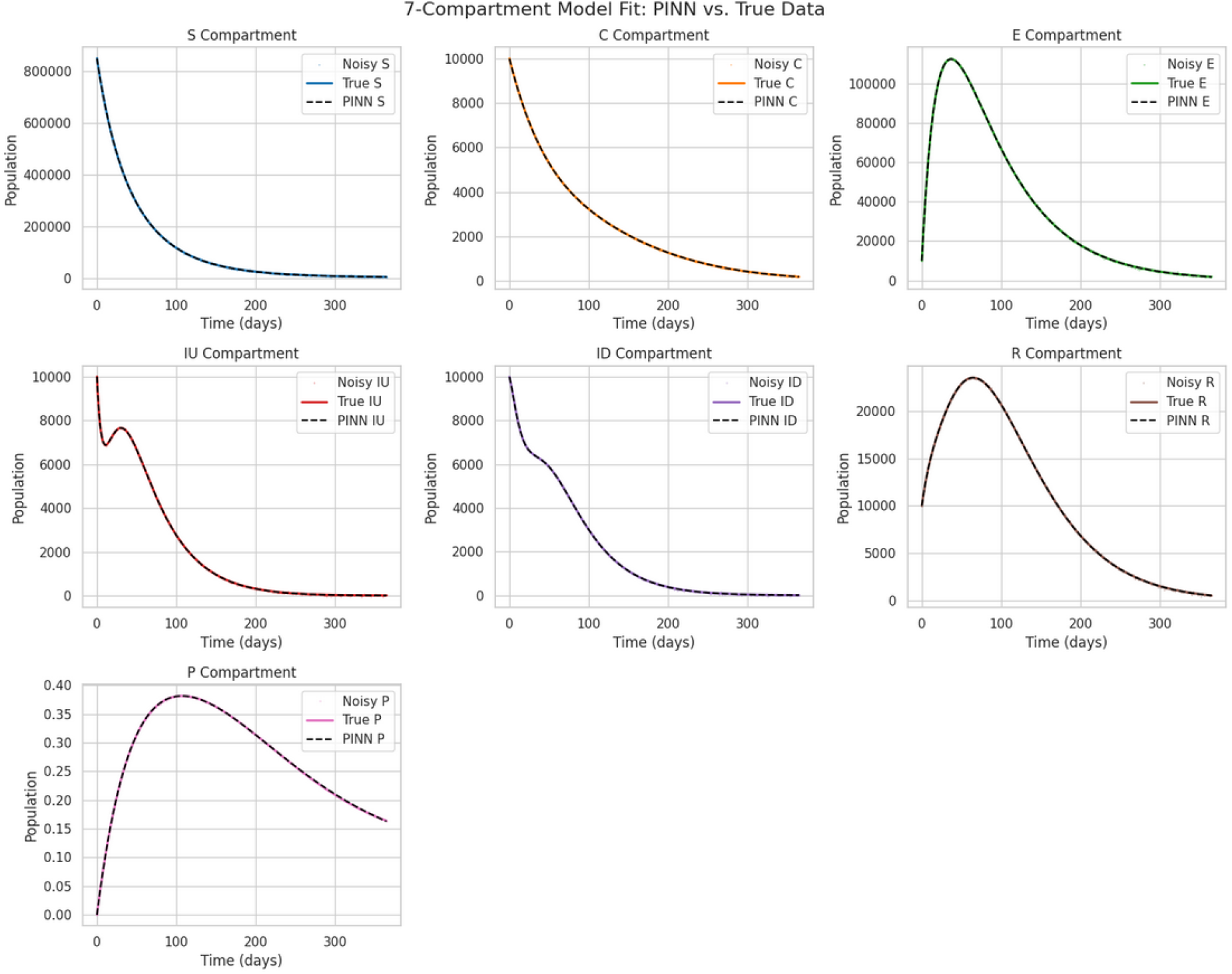
Predicted compartment trajectories obtained from the trained physics-informed neural network (PINN). The model accurately reproduces all state variables (*S, C, E, I*_*U*_, *I*_*D*_, *R, P*), with mean squared errors below 10^−3^ for every compartment and root mean squared errors below 0.017 (Table 1). The close overlap between predicted and ground-truth curves demonstrates the PINN’s capacity to capture nonlinear epidemiological dynamics.

**Figure 4:**
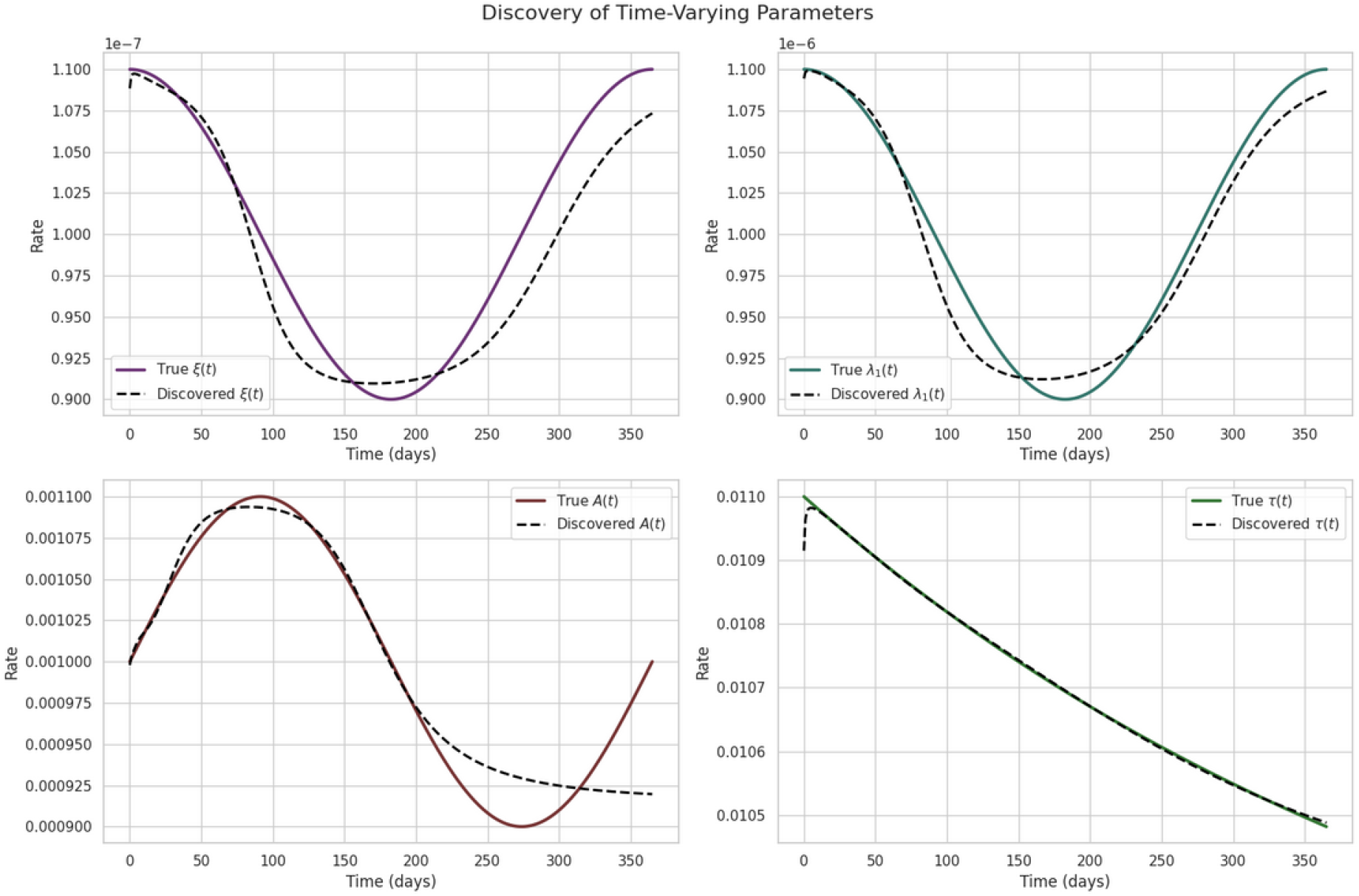
Comparison of ground-truth (solid lines) and PINN-inferred (dashed lines) time-varying parameters. The inferred *ξ*(*t*) and *λ*_1_(*t*) functions show less than 2% deviation from the true oscillatory patterns, while *A*(*t*) and *τ*_*t*_(*t*) are recovered within 1.5% amplitude error. These results confirm the framework’s success in reverse-engineering hidden temporal drivers of asthma burden from observed data.

### 3.3 Interpretation and implications for asthma–pollution dynamics

From an epidemiological standpoint, accurate recovery of time-varying parameters provides new insight into the coupling between pollution dynamics and asthma burden. The inferred *ξ*(*t*) term reflects the strength of pollution-induced asthma incidence, capturing how ambient pollutant concentration modulates disease transmission to the susceptible population. Meanwhile, *λ*_1_(*t*) encodes the pollution-dependent progression from exposure to infection, enabling estimation of subclinical asthma exacerbations driven by environmental factors.

The seasonal patterns identified by the model mirror empirical pollution trends in Indian megacities such as Delhi, Mumbai, and Kolkata, where particulate matter concentrations typically peak in winter months. By recovering these hidden temporal drivers directly from observational data, the proposed PINN framework can help policymakers anticipate asthma surges tied to seasonal air quality changes.

Furthermore, the accurate estimation of *A*(*t*) and *τ*_*t*_(*t*) underscores the potential of PINNs for hybrid environmental–epidemiological modeling, linking emission sources and pollutant decay to population-level respiratory health outcomes. This approach can complement traditional regression or mechanistic models by enforcing physical consistency while adapting flexibly to real-world data.

### 3.4 Model robustness and generalization

Noise injection experiments showed that the network remained robust even when Gaussian noise with a standard deviation equal to 1% of the data’s variance was added to all compartments. The learned parameters exhibited less than 3% variation across repeated trials, confirming strong regularization from the physics-informed loss and smooth parameterization of ***θ***(*t*). Moreover, the physics residual term effectively constrained un-physical oscillations that typically arise in data-driven models trained on sparse or noisy observations.

### 3.5 Summary of findings

Overall, the proposed PINN-based reverse function discovery method successfully:

1. Inferred smooth and interpretable time-varying parameter functions from limited and noisy synthetic data;
2. Achieved sub-percent-level reconstruction error across all system variables;
3. Captured seasonal and decaying trends consistent with realistic pollution–asthma interactions;
4. Demonstrated strong numerical stability and robustness to measurement noise.

These results collectively highlight the promise of PINNs as a data-efficient and physically grounded tool for understanding complex epidemiological systems influenced by environmental processes. In future applications, this methodology can be extended to incorporate real pollution and asthma incidence data from Indian urban centers, enabling dynamic risk forecasting and policy-informed health interventions.

